# Opioids for breathlessness: Psychological and neural factors influencing response variability

**DOI:** 10.1101/344523

**Authors:** Sara J. Abdallah, Olivia K. Faull, Vishvarani Wanigasekera, Sarah L. Finnegan, Dennis Jensen, Kyle Pattinson

## Abstract

Effective management of distressing bodily symptoms (such as pain and breathlessness) is an important clinical goal. However, extensive variability exists in both symptom perception and treatment response. One theory for understanding variability in bodily perception is the ‘Bayesian Brian Hypothesis’, whereby symptoms may result from the combination of sensory inputs and prior expectations. In light of this theory, we explored the relationships between opioid responsiveness, behavioural/physiological symptom modulators and brain activity during anticipation of breathlessness.

**Methods:** We utilised two existing opioid datasets to investigate the relationship between opioid efficacy and physiological/behavioural qualities, employing hierarchical cluster analyses in: 1) a clinical study in chronic obstructive pulmonary disease, and 2) a functional magnetic resonance brain imaging study of breathlessness in healthy volunteers. We also investigated how opioid efficacy relates to anticipatory brain activity using linear regression in the healthy volunteers.

**Results:** Consistent across both datasets, diminished opioid efficacy was more closely associated with negative affect than with other physiological and behavioural properties. Furthermore, in healthy individuals, brain activity in the anterior cingulate and ventromedial prefrontal cortices during anticipation of breathlessness were inversely correlated with opioid effectiveness.

**Conclusion:** Diminished opioid efficacy for relief of breathlessness may be associated with high negative affective qualities, and was correlated with the magnitude of brain activity during anticipation of breathlessness.

**Clinical implications:** Negative affect and symptom expectations may influence perceptual systems to become less responsive to opioid therapy.

## INTRODUCTION

Chronic breathlessness is a multidimensional and aversive symptom, which is often poorly explained by underlying pathophysiology (1). For many patients, breathlessness is refractory to maximal medical therapies targeting disease processes (2). However, opioids are thought to be a possible therapeutic avenue to treat symptomology independently of disease (3, 4). Importantly, research in other aversive symptoms (such as chronic pain) has demonstrated that qualities such as anxiety and depression (collectively termed *negative affect* here) can both exacerbate symptoms (5) and reduce opioid efficacy (6, 7). Therefore, it may be pertinent to consider such behavioural factors when contemplating the use of opioids for breathlessness.

According to the Bayesian Brain Hypothesis, perception (e.g., breathlessness) is the result of a delicate balance between the brain’s set of expectations and beliefs (collectively known as *priors*), and incoming sensory information (8, 9). An individual’s priors are shaped by previous experiences and learned behaviours. For example, if climbing a flight of stairs triggers severe breathlessness, an individual may “expect” to experience severe breathlessness during subsequent stair climbing. Negative affect may act as a moderator within this perceptual system (8-11), altering the balance between priors and sensory inputs to influence and potentially exacerbate symptom perception.

In a recent clinical study, Abdallah et al. (4) demonstrated that 11 of 20 (55%) adults with advanced chronic obstructive pulmonary disease (COPD) reported clinically significant relief (defined as a decrease by ≥ 1 Borg units) of exertional breathlessness following single-dose administration of immediate-release oral morphine. While the authors were unable to elucidate the physiological mechanisms underlying opioid response variability, they speculated that unmeasured differences in “*conditioned anticipatory/associative learning*” might play a role.

Therefore, the aim of the present study was to better understand the mechanisms underlying variability in opioid responsiveness for the relief of breathlessness. We have reanalysed data from 1) Abdallah et al. (4) and 2) a behavioural and functional neuroimaging dataset in healthy volunteers by Hayen et al. (12), where the perception of laboratory-induced breathlessness was manipulated with the opioid remifentanil. As with Abdallah et al. (4), Hayen et al. (12) observed variability in responsiveness to opioid therapy, with 9 of 19 (47%) subjects reporting a remifentanil-induced decrease in experimentally-induced breathlessness by ≥10 mm on a 100-mm visual analogue scale (VAS). This parallel approach allowed us to verify associations observed in a clinical population in an independent sample that were free of the confounds of chronic disease, and to shed light on associated changes in brain activity with opioid efficacy. Furthermore, the healthy volunteer dataset allowed us to test the hypotheses presented in Abdallah et al. (4) regarding how potential differences in conditioned anticipatory/associative learning might influence variability in the effect of opioids upon breathlessness.

## METHODS

A brief overview of the study methods is provided here. Full methodological details can be found in the supplementary material found in pages 21 to 43.

### COPD exercise study

Twenty participants with severe COPD (mean±SD forced expiratory volume in 1 sec % predicted (FEV_1_%predicted): 35±9) completed two sessions (morphine 0.1 mg/kg or placebo – randomized order and double-blinded), where physiological and perceptual parameters were measured during constant-load cardiopulmonary cycle exercise testing at 75% of peak incremental cycle power output. Intensity and unpleasantness of breathlessness were rated using Borg’s modified 0-10 category ratio scale at rest and during exercise (13). Specifically, subjects were asked “How intense is your sensation of breathing overall?” and “How unpleasant or bad does your breathing make you feel?” (see online data supplement for details). Data were analyzed at isotime, defined as the highest equivalent 2-min interval of exercise completed by a given participant during constant-load cardiopulmonary cycle exercise tests performed after treatment with oral morphine and placebo. The change in all scores were calculated as opioid minus placebo. Participants were also characterized using questionnaires listed in **Table 1.**

**Table 1.**
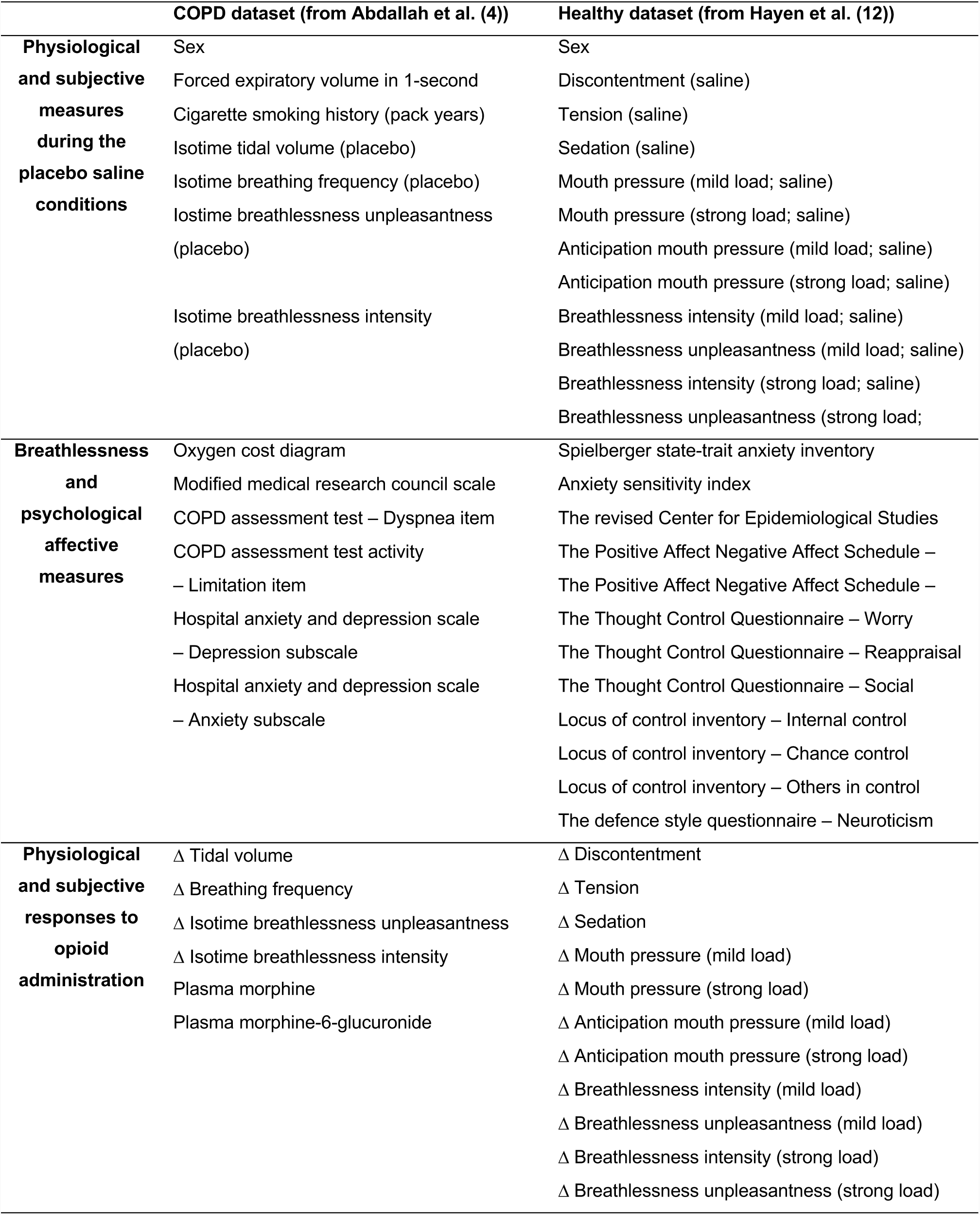
Variables included in the hierarchical cluster analysis.

### FMRI study in healthy volunteers

In the original study (12), 19 healthy participants underwent two functional magnetic resonance imaging (FMRI) scans (3T Siemens Trio scanner), on two separate visits. An intravenous infusion of either remifentanil or saline placebo was delivered in a counterbalanced, randomised and double-blind fashion. The remifentanil dose was modelled on a concentration of 0.7 ng/ml in the brain, achieved using a target-controlled infusion pump. Experimental breathlessness was induced using inspiratory resistive loading combined with mild hypercapnia. Participants also underwent a delay-conditioning paradigm before the scanning visits, wherein they learned associations between three visual cues (shapes) presented on a screen, and three conditions: mild inspiratory load (approximately −3 cmH_2_0), strong load (approximately −12 cmH_2_0) and unloaded breathing. A cued anticipation period of 8 seconds then preceded each loading condition. Participants rated the intensity and unpleasantness of their breathlessness using a VAS (0-100 mm). The change in all scores was calculated as opioid minus placebo. Participants were also characterized using questionnaires listed in **Table 1.**

During the breathlessness protocol, a familiarisation phase of at least 5 minutes was first allowed for each participant to become accustomed to the breathing system. Throughout the experiment, the end-tidal partial pressures of carbon dioxide and oxygen were maintained constant by manual adjustment of inspiratory gasses.

### Hierarchical cluster analyses

Studies in chronic pain have demonstrated an inverse relationship between negative affect and opioid analgesia (6, 7). Therefore, we explored the possible relationships between the magnitude of opioid-induced relief of breathlessness, behavioural measures and physiological traits. A hierarchical cluster analysis (MATLAB: 2013a, MathWorks Inc., Natick, MA, USA) was performed on questionnaires, breathlessness ratings, and physiological measures from each of the two datasets (**Table 1**). Despite differences in the experimental protocols of Abdallah et al. (4) and Hayen et al. (12), the measurement tools used in both studies broadly represented similar domains (see **Table 1)**.

Variables were first aligned such that larger values represented more negative properties. This was achieved through the multiplication of relevant variables by −1 (e.g., FEV_1_% predicted was multiplied by −1 as a larger FEV_1_%predicted value reflects less severe disease). All measures were then individually normalised via Z-transformation, to allow accurate variable comparisons and distance calculations.

Hierarchical cluster models reorder variables based on their correlation strengths, such that groups of related measures sit closer to each other than non-related measures. This modeling process formalizes the relationship between pairs of variables and the manner by which shared variance can be described as part of larger, related clusters. The clustering algorithm is a form of unsupervised machine learning that initially considers pairs of variables in terms of their similarity or “distance”, and will therefore find links between groups of measures regardless of their significance. Linked pairs are incorporated into larger clusters with the goal of minimizing a cost function (distance to be bridged) - a process that can be thought of as minimizing the dissimilarity within clusters. As pairs become clusters, a cluster tree or dendrogram is created. The distance between neighboring branches indicates the relative similarity of two measures. The branching of these levels is an arbitrary distance measure and does not indicate the strength of the correlation in terms of a Pearson’s R value.

A ‘threshold of relatedness’ can then be applied to the hierarchical cluster tree, to formally separate clusters of variables thought to be statistically distinct from one-another. In applying this ‘threshold of relatedness’, the most mathematically distinct cut point needs to be considered together with the most biologically relevant information, given an a-priori hypothesis. We applied a typical threshold technique of the ‘elbow method’ to identify the most mathematically distinct clusters, before considering further cluster divisions utilizing a-priori knowledge and visual inspection of the dendrogram structure. This analysis allowed us to explore natural ‘groupings’ within the recorded measures, and the relationship of these groupings to each other. Clusters were defined by higher-order variable groupings denoted by the hierarchical clustering structure, with a minimum intra-cluster correlation coefficient of 0.3 between the variables.

### FMRI analysis of opioid efficacy

In the present study, the FMRI data from the healthy volunteers was then investigated in consideration of the Bayesian Brain Hypothesis. We explored how brain activity associated with anticipation of breathlessness (during the saline placebo condition) may relate to an individual’s ‘opioid efficacy’ for the treatment of breathlessness. The group of items that formed **Cluster B** within the hierarchical cluster analysis on the healthy volunteers (see **Fig. 1**) were used to define overall ‘opioid efficacy’ (i.e., items that represented opioid-induced changes in physiological and subjective measures). These items included opioid-induced changes in: discontentment, tension and mouth pressure during anticipation of mild and strong loading; mouth pressure and breathlessness intensity and unpleasantness during mild and strong loading; and sedation during the saline condition. We undertook the data reduction technique of a principal component analysis (PCA; MATLAB 2013a) on this group of variables, and the resulting individual scores were included within a group FMRI analysis of the saline placebo condition only, using a general linear model (*Z* > 2.3, whole brain corrected *p* < 0.05).

**Figure 1.**
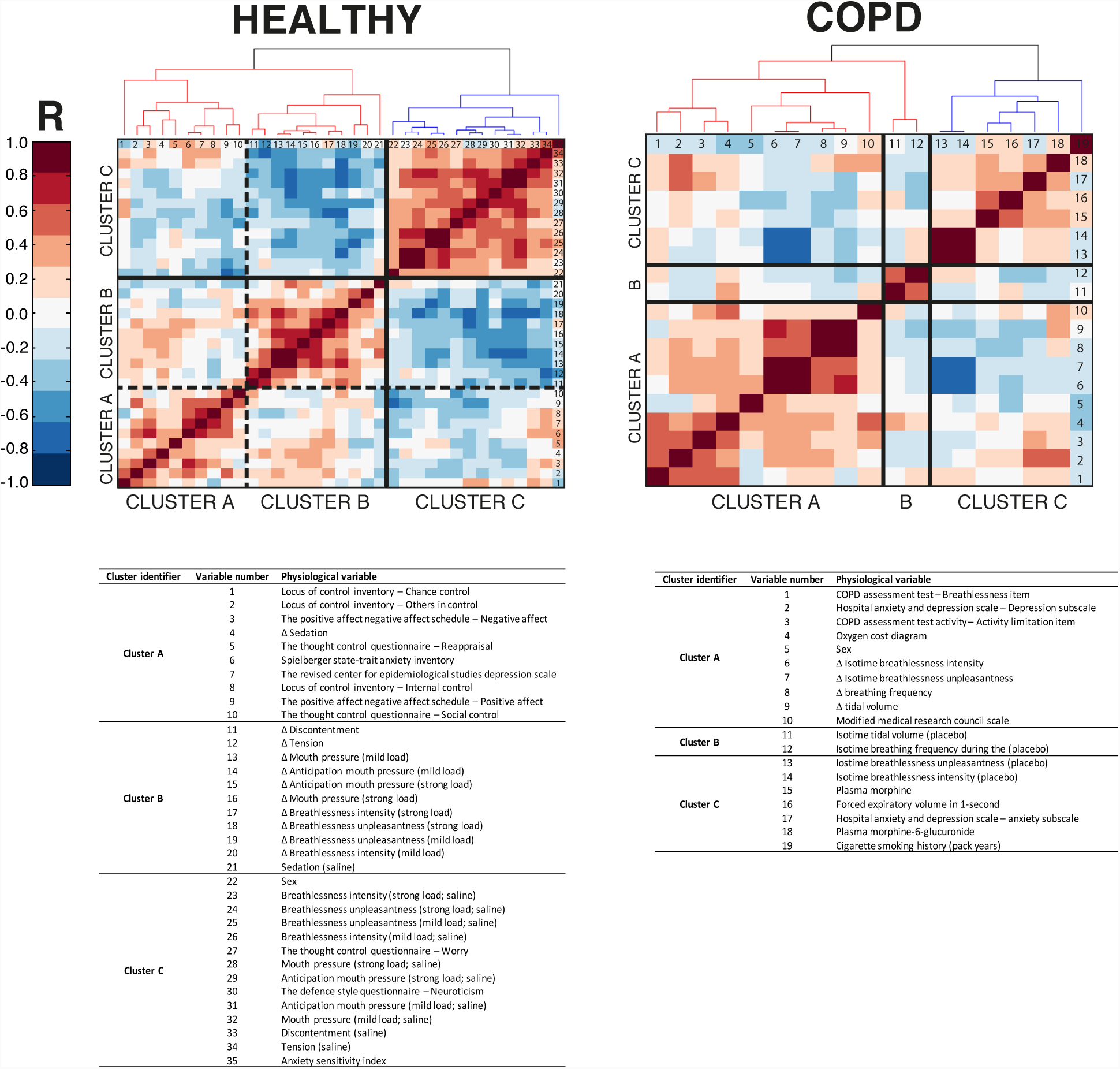
Clustergram of physiological and behavioural variables in healthy volunteers and participants with COPD (top panel). Identified hard cluster boundaries (via ‘elbow method’) are denoted in solid black lines, whilst sub-clusters (via visual inspection) are denoted with dashed lines. Tables identify the physiological and behavioural variables induced in each of the sub-clusters (bottom panel). The change (Δ) in all scores was calculated as: opioid minus placebo. In the COPD dataset, physiological and perceptual responses were evaluated during exercise at isotime - defined as the highest equivalent 2-min interval of exercise completed by each participant after oral morphine and placebo.

## RESULTS

The breathlessness stimuli in the COPD (exercise) and healthy volunteer (inspiratory resistive load) datasets both increased perceptions of work-effort and air hunger during the saline condition (see original manuscripts (4, 12) and **Supplementary Fig. 5**). In both groups, there was significant inter-individual variability in the magnitude of opioid-induced relief of breathlessness.

### Hierarchical cluster analysis

The hierarchical cluster analysis of the COPD group supported the existence of three distinct clusters of variables, which were verified by the elbow method (**Fig. 1,** and **Supplementary Fig. 4**).

In the COPD dataset (**Fig. 1**), 4/10 items in **Cluster A** represented physiological and subjective responses to opioid administration, and 5/10 items represented breathlessness and psychological affective measures. This **‘response-affect’** cluster included the: COPD assessment test (CAT) breathlessness and activity-limitation items (14); hospital anxiety and depression scale (HADS) - depression subscale (15); oxygen cost diagram (16); modified medical research council dyspnoea scale (mMRC) (17); opioid-induced changes in isotime breathlessness intensity and unpleasantness, and breathing frequency and tidal volume. **Cluster B** consisted of 2 items that represent baseline (placebo condition) physiological measures: isotime tidal volume and breathing frequency. **Cluster C** consisted of 7 items, 4 of which represented baseline physiological and subjective measures at rest and during the placebo condition. This **‘baseline’** cluster included: breathlessness intensity and unpleasantness at isotime during the placebo condition; cigarette smoking history; FEV1%predicted; plasma morphine concentrations; plasma morphine-6-glucuronide concentrations; and the HADS anxiety subscale (15).

In the healthy volunteer dataset, the elbow method initially supported the existence of two distinct clusters (**Fig. 1**, solid lines; and **Supplementary Fig. 4**). Upon visual inspection, the larger cluster could clearly split further into two distinct and related clusters (**Fig. 1**, dashed lines). **Cluster A** appeared to be a predominantly **‘affective’** cluster, consisting of 10 items - 9 of which are representative of psychological affective measures. These were the Levenson Multidimensional Locus of Control Inventory– Chance Control, Others in control, and Internal control subscales (18), The thought control questionnaire - Social Control, and Reappraisal subscales (19); Positive and Negative affect schedules (20); the Spielberger State-Trait Anxiety Inventory (21); the revised Center for Epidemiological Studies Depression Scale (22). The remaining item in this cluster is opioid-induced change in sedation (23). **Cluster B** was representative of physiological and subjective responses to opioid administration, as 9/10 items in this cluster were related to opioid-induced changes (see **Fig. 1** – **Cluster B** list of the healthy dataset). The remaining item in this cluster was baseline sedation in the saline placebo condition. **Cluster C** predominantly represented (11/14 items) physiological and subjective measures during the baseline (saline placebo) condition. These items were: breathlessness intensity and unpleasantness in the strong and mild load; mouth pressure during anticipation of breathlessness in both strong and mild loading; discontentment and tension (23) during the saline placebo condition; and sex. The remaining 3 items were related to the affective domain; the Thought Control Questionnaire – Worry subscale (19); the Defence Style Questionnaire – Neuroticism subscale (24); and Anxiety sensitivity index (25).

In both datasets, the predominant ‘affect’ and ‘response’ clusters were more closely related to each other than to the ‘baseline’ cluster. Importantly, the association (smaller distances between respective clusters) between the ‘affect’ and ‘response’ clusters indicated that worse affective scores corresponded to a smaller degree of opioid-induced relief of breathlessness (i.e. variables were aligned such that larger values represented more negative properties).

### FMRI analysis

The FMRI analysis revealed significant *anticipatory* brain activity that directly correlated with opioid unresponsiveness (i.e., greater brain activity correlated with a smaller ‘response’ score from the PCA of **Cluster B** - the sub-cluster representing ‘opioid responsiveness’) in the anterior cingulate cortex (ACC) and ventromedial prefrontal cortex (vmPFC) during both mild and strong loading; and the caudate nucleus (CN) during anticipation of mild loading only (**Fig. 2**). That is, the greater the activity in these brain regions during anticipation of breathlessness, the smaller the degree of opioid-induced relief of breathlessness.

**Figure 2.**
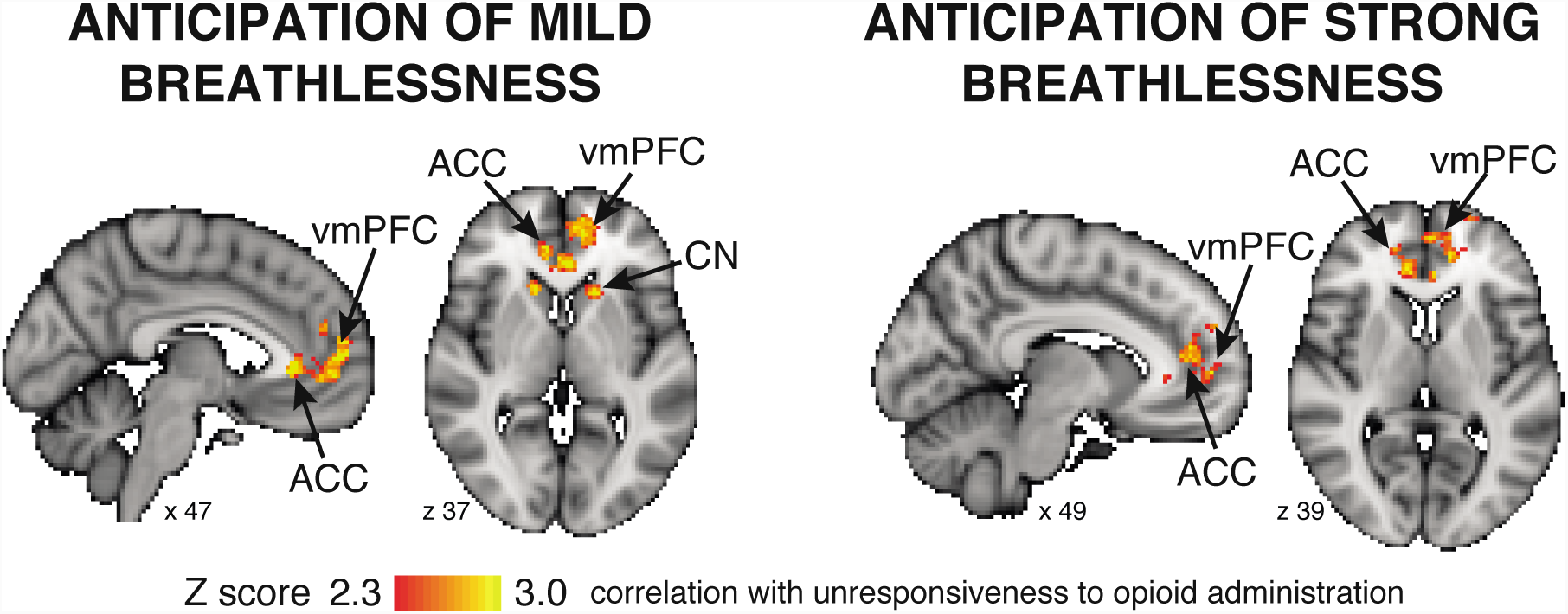
Mean BOLD changes identified during anticipation of the mild (left panel) and strong breathlessness (right panel) challenge. The image consists of a colour-rendered statistical map superimposed on a standard (MNI 2×2×2) brain. Significant regions are displayed with a threshold Z > 2.3, using a cluster probability threshold of p < 0.05. ACC, anterior cingulate cortex; CN, caudate nucleus; vmPFC, ventromedial prefrontal cortex.

## DISCUSSION

In this study, we have shown that diminished opioid efficacy for breathlessness was more closely associated with negative affect than other physiological and behavioural properties in both a COPD and a healthy volunteer dataset. Furthermore, functional neuroimaging findings of the healthy volunteer study revealed that the magnitude of opioid-induced relief of breathlessness was inversely correlated with brain activity in the ACC and vmPFC during anticipation of breathlessness, under a saline placebo infusion. These findings suggest that opioid efficacy for breathlessness may be associated with broader negative affective qualities within an individual and may be directly related to brain activity during conditioned responses to breathlessness stimuli.

Our findings from the behavioral data in both studies suggest that opioid responsiveness is inversely associated with the collective co-existing weight of affective moderators (**Fig. 1**). That is, individuals with high vs. low negative affect are less likely to experience opioid-induced relief of breathlessness. These results align with previous work in chronic pain, where it has been found that in addition to less effective analgesia, negative affective qualities are associated with dose escalation (26) and greater difficulty in reducing opioid medication use (27). Interestingly, the hierarchical cluster analysis revealed a ‘baseline’ cluster in COPD dataset that was not as tightly correlated as the healthy dataset. These results support the view that in COPD, breathlessness is a multifactorial symptom that is likely influenced by variables other than baseline pulmonary function. Notably, we have demonstrated that opioid responsiveness was not related to plasma levels of morphine, as plasma morphine and plasma morphine-6-glucuronide levels in the COPD dataset were more closely related to physiological measures at baseline than to opioid responsiveness. These findings are in keeping with the results of Abdallah et al. (4) who found no correlation between plasma morphine and morphine-6-glucuronide levels and opioid-induced changes in exertional breathlessness.

Interestingly, the cluster structure revealed in the COPD participants was conceptually consistent with that found in the healthy volunteers. Free of the confounds of respiratory disease and chronic breathlessness, the results in these healthy individuals suggest that even subtle variations in affective traits may have measurable effects upon opioid responsiveness. Our results suggest that individual affective traits, and not baseline levels of breathlessness *per se*, may contribute towards the magnitude of opioid responsiveness.

The healthy volunteer dataset also allowed us to examine the relationship between opioid efficacy and anticipatory brain activity in the context of the Bayesian Brain Hypothesis. Opioid-induced changes (quantified by the scores form the ‘response’ cluster variables) were directly related to *anticipatory* brain activity towards an upcoming breathlessness stimulus under baseline (placebo) conditions. The activated regions - ACC and vmPFC - are contributors to the brain network involved in evaluating the relevance of a stimulus and its associated value (i.e., reward processing), and are thought to be involved in generating predictions on emotional state and bodily awareness (9, 28). When anticipating breathlessness, individuals with greater brain activity in these regions were less likely to experience meaningful opioid-induced relief of breathlessness, and therefore potentially more ‘resistant’ to opioid therapy. In particular, associated negative affective properties might influence breathlessness perception by more heavily weighting the brain’s perceptual system towards learned associations (or priors) during anticipation of breathlessness (8). For example, in anticipation of climbing a set of stairs, an individual with worse affective traits may have greater breathlessness expectations (e.g., via ‘catastrophizing’ the severity of breathlessness during previous experiences of stair climbing) relative to an individual with more ‘normal’ affective traits. In turn, and despite receiving the same sensory afferent inputs when they do climb the stairs, the individual with worse affective traits may be less responsive to opioid therapy as their breathlessness perception is more rigidly attracted towards their breathlessness expectations (i.e., strong priors) relative to the individual with ‘normal’ affective traits.

Finally, whilst this neuroimaging work was completed in healthy volunteers, previous neuroimaging studies have evaluated the relationship between learned associations and relief of breathlessness in COPD. In a study of pulmonary rehabilitation, and in contrast to the present findings with opioids, Herigstad et al. (29) reported that baseline activity in the brain network responsible for generating predictions (e.g., ACC) correlated positively with changes in breathlessness following pulmonary rehabilitation in COPD. Pulmonary rehabilitation is thought to exert its benefits, in part, by re-shaping associations and modulating negative affect (29-31). The results of these studies suggest that individuals with strong learned associations (priors) and negative affective comorbidities may be more likely to benefit from treatments such as pulmonary rehabilitation than opioids for relief of breathlessness. It is also possible that individuals with these strong associations and negative affective comorbidities may require higher opioid doses to experience adequate relief of breathlessness, as previously demonstrated in pain (6, 26, 27).

### Methodological considerations

Interpretation of the cluster analysis is limited by sample size, which influences model sensitivity and stability. Given more subjects, techniques such as factor analysis could provide additional statistical confidence. Factor analysis allows a formal model to be fitted to the data. This allows investigation of the relationship between variables and establishes whether an underlying shared construct exists, via a description of the full variance of the data and a number of fit statistics. Although we identified several clear clusters that would easily feed into a confirmatory factor analysis, it is generally accepted that for this technique the number of samples should be at least 5 times greater than the number of investigatory variables (32). This would limit us to a model consisting of 4 variables, which we believe does not accurately describe the clusters identified in this dataset. Therefore, we selected the elbow method as a cluster threshold technique, which proposed 2 or 3 clusters for the healthy population and 3 clusters for the COPD population; these clusters were validated and refined visually.

A second important consideration of hierarchical cluster analysis is that omitted variables cannot be inferred upon, and that unrelated variables may randomly cluster together. Despite these limitations, our ability to replicate the behavioural results across two independent datasets increases confidence in our findings. Nevertheless, further replications with larger sample sizes will allow more thorough investigation into the major contributing affective variables. For example, in contrast to the healthy dataset, the COPD dataset measured a smaller number of psychological affective measures. In addition to HADS depression, the “response-affect” cluster in the COPD dataset included the mMRC, oxygen cost diagram and the CAT breathlessness item. This clustering may be indicative of the fact that the mMRC, oxygen cost diagram and CAT are associated with anxiety and depression in COPD (33, 34). Alternatively, this may relate to the fact that in this small sample size, the hierarchical cluster model may be less stable than when compared to a larger sample size.

## Conclusions

In conclusion, the datasets by Abdallah et al. (4) and Hayen et al. (12) have allowed us to explore predictors of opioid responsiveness, and to generate hypotheses based upon potential neurobiological mechanisms of action. Although additional research is necessary, our results are novel and support the hypothesis that opioids may be less effective for the treatment of breathlessness among individuals with higher levels of negative affect comorbidities and strong learned associations. Our results provide clues towards opioid mechanisms of action for relief of breathlessness, which could be tested in future prospective and longitudinal studies, especially as we move towards individualized, safe and targeted symptom management.

## Acknowledgments

The authors wish to thank the contributions of Dr Anja Hayen, Dr Mari Herigstad, Steward Campbell, Payashi Garry, Simon Raby, Josephine Robertson, and Ruth Webster towards the data collection for the healthy volunteer study.

## SUPPLEMENTARY MATERIAL

## INTRODUCTION

The original study by Abdallah et al. (1) investigated 1) the efficacy of oral morphine in reducing exertional breathlessness and improving exercise tolerance in patients with advanced chronic obstructive pulmonary disease (COPD), and 2) the physiological (e.g. minute ventilation) and perceptual (e.g. ratings of breathlessness at peak exercise during incremental cycle exercise testing) factors mediating opioid responsiveness. Meanwhile, the study by Hayen et al. (2) explored functional brain activity during perception of conditioned mild and strong breathlessness stimuli, following either saline placebo or remifentanil infusion in healthy participants.

Abdallah et al. (1) induced breathlessness in adults with COPD using a constant-load cycle exercise test. Hayen et al. (2) induced breathlessness in healthy adults using an inspiratory resistive load combined with mild hypercapnia (details provided below). The stimuli selected in both studies increased breathing intensity and unpleasantness during the saline condition. In both studies, opioid decreased breathlessness intensity and unpleasantness. Importantly, significant inter-individual variability in responsiveness to opioid therapy was demonstrated in both studies. In Abdallah et al. (1), 11 out of 20 adults with COPD reported a morphine-induced decrease in breathlessness intensity that met or exceeded the minimally clinically important difference (MCID) of 1 Borg unit (3). In Hayen et al. (2), 9 out of 19 healthy adults reported a remifentanil-induced decrease in breathlessness intensity that met or exceeded 10 mm on 100 mm visual analogue scale (VAS) (3, 4). The mechanisms mediating this variability in responsiveness to opioid therapy are currently unknown.

In the current study we have undertaken a reanalysis of behavioural and physiological measures collected by Abdallah et al. (1) and Hayen et al. (2) (including previously unpublished data) to investigate the association between behavioural measures and opioid responsiveness in both health and COPD, and second, to identify brain regions that may contribute to this relationship. These questions regarding inter-individual variability were not the primary aims of the two original studies.

## METHODS

### COPD study

#### Participants

Participants included men and women aged ≥40 yrs with clinically stable Global Initiative for Obstructive Lung Disease stage 3 or 4 COPD (5) and chronic breathlessness. This was defined as a modified Medical Research Council dyspnoea score of ≥3 (6), a Baseline Dyspnoea Index focal score of ≤6 (7) and/or an Oxygen Cost Diagram rating of ≤50% full scale (8), despite optimal treatment with bronchodilators, corticosteroids and/or phosphodiesterase inhibitors (5). Exclusion criteria included: smoking history <20 pack-years; change in medication dosage and/or frequency of administration in the preceding 2-weeks; exacerbation and/or hospitalization in preceding s6-weeks; arterialized capillary PCO_2_ (P_ac_CO_2_) >50 mmHg at rest; presence of other medical condition(s) that could contribute to breathlessness and/or exercise intolerance; important contraindications to cardiopulmonary exercise testing (CPET); self-reported history of addiction and/or substance abuse; use of anti-seizure drugs or opioids; use of daytime oxygen; and exercise-induced oxyhemoglobin desaturation to <80% on room air.

#### Study design

This single-center, randomized, double-blind, placebo-controlled, crossover trial (ClinicalTrials.gov identifier NCT01718496) consisted of two intervention periods separated by a washout period of ≥48 hrs. Participants were randomized in a 1:1 ratio to receive immediate-release oral morphine sulphate (0.1 mg/kg body mass; Statex™, Paladin Labs Inc., Montreal, QC, Canada) or diluted simple syrup (placebo) prepared in 250 ml of orange juice. A computer-generated block randomization schedule was prepared by a third-party not involved in the trial. The study protocol and informed consent form received ethical approval from Health Canada (File No. 9427-M1647-48C) and the Research Ethics Board of the Research Institute of the McGill University Health Centre (MP-CUSM-12-325-T).

After providing written informed consent, participants completed a screening/familiarization visit followed by two randomly assigned treatment visits. During Visit 1 participants completed: ***behavioural questionnaires*** including the Hospital Anxiety and Depression Scale (9), the modified Medical Research Council dyspnoea scale (6), the Oxygen Cost Diagram (8) and the COPD Assessment Test (10); post-bronchodilator (400 µg salbutamol) pulmonary function testing; and a symptom-limited incremental cardiopulmonary cycle exercise test (CPET) to determine peak power output, defined as the highest power output that the participant was able to sustain for 30-sec or longer. At the beginning of Visits 2 and 3, participants inhaled 400 µg of salbutamol to standardize the time since last bronchodilator administration. Participants were then administered oral morphine or placebo. Thirty-minutes thereafter, blood was collected for measurements of plasma concentrations of morphine and its metabolites: morphine-3-glucuronide and morphine-6-glucuronide. Participants then completed a symptom-limited constant-load cycle CPET at 75% peak power output.

Symptom-limited exercise tests were conducted on an electronically braked cycle ergometer (Lode Corival, Lode B.V. Medical Tech., Groningen, The Netherlands) using a computerized CPET system (Vmax Encore™ 29C). Incremental CPETs consisted of a steady-state rest period of at least 6-min, followed by 1-min of unloaded pedalling and then 5 W/min increases in power output. Constant-load CPETs consisted of a steady-state rest period of at least 6-min, followed by 1-min of unloaded pedaling and then a step increase in power output to 75% peak power output. Cardiac, metabolic, breathing pattern and gas exchange parameters were collected and analyzed as previously described (11). Using Borg’s modified 0-10 category ratio scale (12), participants rated the intensity and unpleasantness of their breathlessness every 2-min during CPET, and at end-exercise. Subjects were asked “How intense is your sensation of breathing overall” and “How unpleasant or bad does your breathing make you feel”. The “radio analogy” was used to distinguish between intensity and unpleasantness of breathlessness (13). Briefly, the radio analogy makes the distinction between the volume of the music emanating from the radio (i.e. intensity) and the unpleasantness of the music being heard. Subjects were instructed that a loud volume (i.e. high intensity) can be pleasant if their favourite song was playing on the radio, and unpleasant if a song they dislike was playing. Therefore, in rating their breathlessness on the Borg’s scale, individuals were specifically rating breathlessness intensity and unpleasantness. Physiological and perceptual responses were evaluated during exercise at isotime – defined as the highest equivalent 2-min interval of exercise completed by each participant. The change in tidal volume, breathing frequency, and breathlessness ratings at isotime were calculated as: morphine – saline placebo.

#### Hierarchical cluster analysis

A full hierarchical cluster analysis (MATLAB: 2013a, MathWorks Inc., Natick, MA, USA) was performed on the: behavioural questionnaires collected during Visit 1; isotime breathing frequency and tidal volume and breathlessness intensity and unpleasantness ratings during the placebo condition; isotime changes in breathing frequency, tidal volume and breathlessness intensity and unpleasantness ratings; and morphine-induced changes in plasma morphine and morphine-6-glucuronide. Morphine-3-glucuronidel levels were not included in the hierarchical cluster analysis as it is an inactive metabolite. These hierarchical cluster analyses allowed us to explore natural ‘groupings’ within the recorded measures in each dataset, and the relationship of these groupings to each other. Clusters were defined by higher-order variable groupings denoted by the hierarchical clustering structure, with a minimum intra-cluster correlation coefficient of 0.3 between the variables. See page 21 for further methodological details on the hierarchical cluster analysis.

## Healthy volunteer FMRI study

This double-blind, randomized, placebo-controlled mechanistic study investigated the neural correlates of the opioid remifentanil upon the anticipation and perception of breathlessness. An aversive delay-conditioning session was followed by two FMRI sessions (remifentanil or saline placebo, counterbalanced across participants). The sessions were performed on three consecutive days at the same time each day.

### Participants

Data from 19 healthy participants (10 females, age 24 (±7 SD) years) was analysed in this study. Written informed consent was obtained in accordance with the Oxfordshire Research Ethics Committee. Although 29 participants originally participated, 10 were excluded for the following reasons: 2 participants exhibited vasovagal syncope during cannulation; 1 participant did not comply with study instructions; 4 participants did not learn the association between visual cues and respiratory stimuli; 3 participants were excluded because of technical difficulties with the MRI equipment. Participants were right-handed non-smokers and had no history of neurological (including painful conditions), pulmonary or cardiovascular disease, were free from acute respiratory infections and were currently not receiving any medication. Participants fasted solids for 6 hours and liquids for 2 hours before every session.

### Initial session

The Center for Epidemiologic Studies Depression Scale (CES-D; (14)) was used to identify (and exclude) participants with clinical depression. The trait scale of the Spielberger State-Trait Anxiety Inventory (STAI) (15) was used to characterize general participant anxiety. The following questionnaires were also collected (these were not analysed in the original manuscript of Hayen et al. (2)): The Anxiety Sensitivity Index (ASI; (16)), the Positive Affect Negative Affect Schedule (PANAS; (17)), the Thought Control Questionnaire (TCQ; (18)), the Defence Style Questionnaire (DSQ-40; (19)), and the Levenson Multidimensional Locus of Control Inventory (LOC; (20)).

### Hierarchical cluster analysis

As with the COPD data set, a full hierarchical cluster analysis was performed on the behavioural questionnaires, mouth pressure and subjective breathlessness scores during the placebo condition; and the change in mouth pressure, subjective breathlessness scores, tension-relaxation and discontentment scores for each level of loading. See page 21 for further details pertaining to the cluster analysis techniques.

### Breathlessness stimulus

The breathlessness stimulus used in this study was intermittent resistive inspiratory loading for 30 to 60 seconds, administered via the magnetic resonance imaging (MRI) compatible breathing system illustrated in **Supplementary Fig. 1**. Manually operated hydraulic valves diverted inspiratory flow via one of three routes that either did not restrict breathing, or provided a mild or strong resistive load. Expiration was unrestricted via a one-way valve (Hans Rudolph, Shawnee, Kansas, USA). The strong load was induced with a static porous glass disc and the mild load was induced with a static spirometry filter. Using these technique, the loads applied were related to the speed and depth of inspiration, and therefore they adapted to physiological differences between individuals or even between breaths. Throughout the experiment, the actual load produced at the mouth was measured. We found no correlation between the pressure generated and the load reported.

**Supplementary Figure 1.**
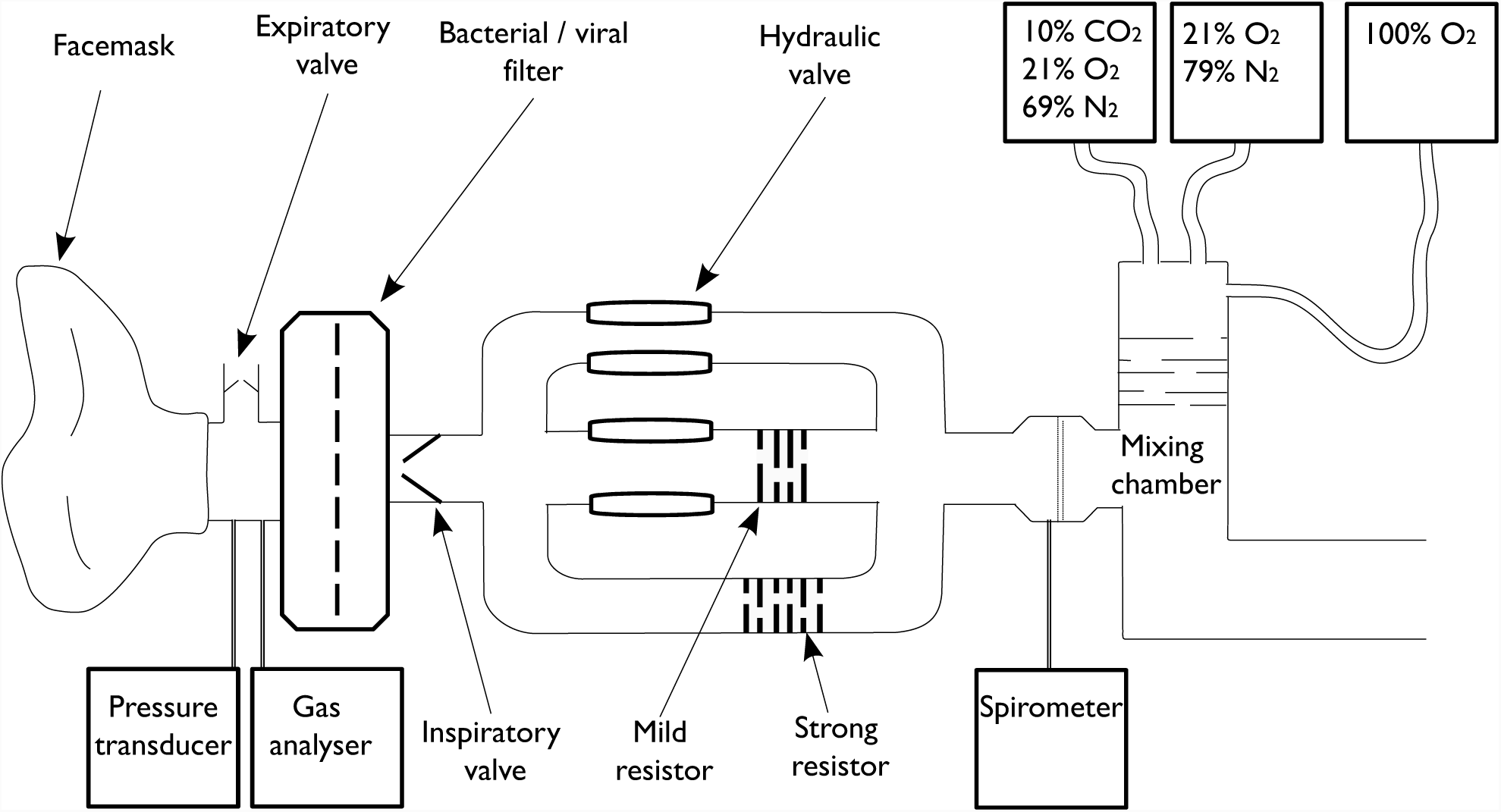
Schematic diagram of respiratory circuit. A facemask (Hans Rudolph, Kansas City, MO, USA) connects to a bacterial and viral filter (Vitalograph, Buckingham, UK) from which respiratory gases and respiratory pressure are sampled via polyethylene extension tubing (Vygon SA, Ecouen, France). One sampling line leads to a pressure transducer (MP 45, ± 50 cmH_2_O, Validyne Corp., Northridge, CA, USA) connected to an amplifier (Pressure transducer indicator, PK Morgan Ltd, Kent, UK). The second sampling line connects to a gas analyzer that samples O_2_ and CO_2_ (ADInstrument Ltd, Oxford, UK). A one-way valve allows expired air to escape close to the mouth in order to minimize rebreathing (Hans Rudolf, Kansas City, MO, USA). The breathing system contains three arms. Participants usually breathe through the first, unobstructed arm. This arm can be closed off by inflating a balloon (embolectomy balloon, Microtek Medical B.V., Zutphen, Netherlands) via a hydraulic system. Closure of the arm forces participants to breathe through the second arm, which contains a pediatric respiratory filter (mild resistor, Intersurgical, Wokingham, UK) and a second balloon valve. Closure of both valves forces breathing through a third arm, which contains a porous glass disk (strong resistor). The three arms recombine into a spirometry module that records respiratory flow (ADInstruments Ltd, Oxford, UK) connected to a custom-made mixing chamber in which medical air, oxygen and 10% CO_2_ in air are combined.

**Supplementary Figure 2.**
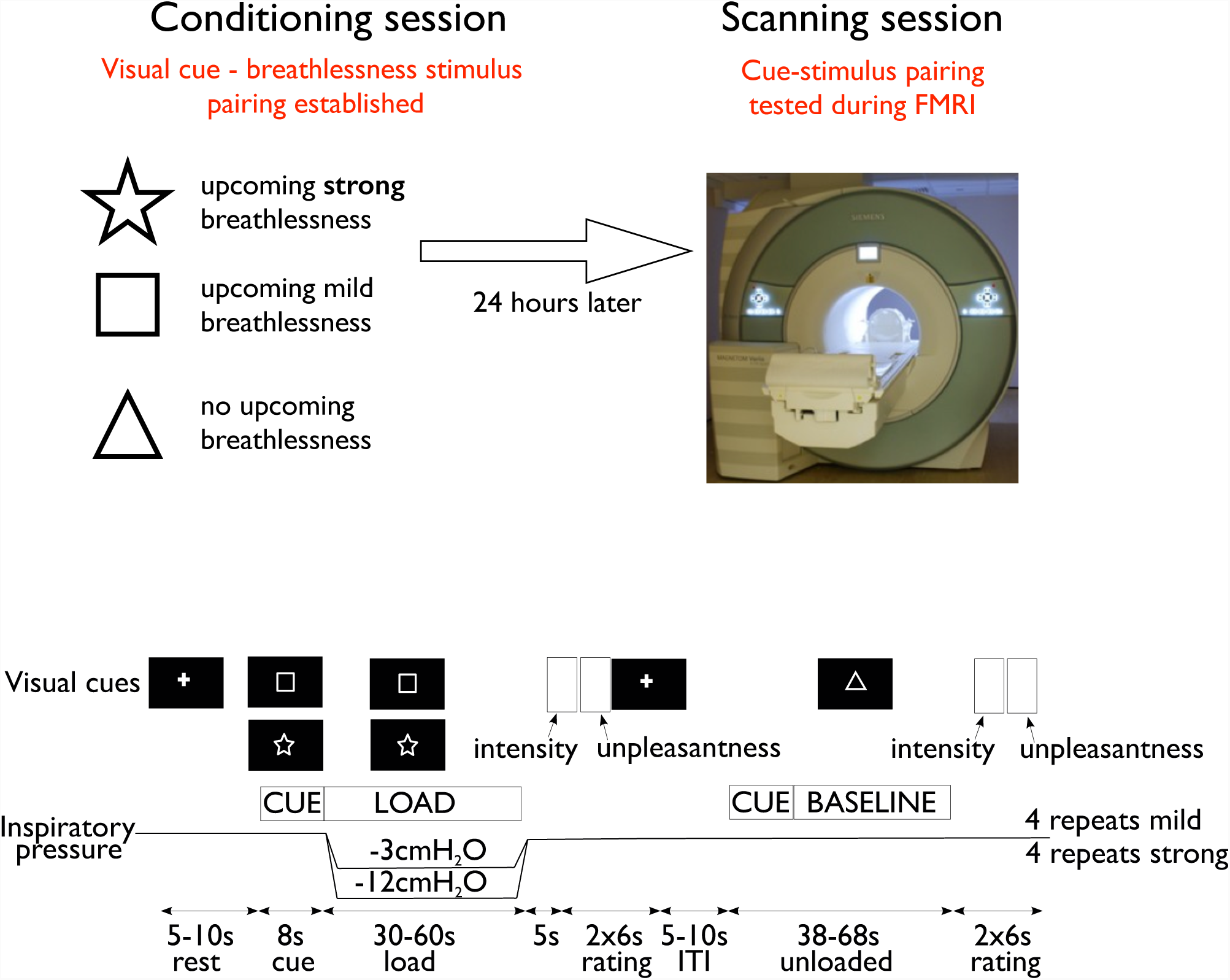
Schematic illustration of experimental session and aversive conditioning paradigm. Prior to the application of each inspiratory load, the fixation cross on the screen changed to one of three shapes, a triangle, a square and a star to signal the imminent application of a stimulus (mild inspiratory load, strong inspiratory load) for eight seconds (anticipation period). The shape remained on the screen during the application of the stimulus (stimulus period) for 30-60seconds and changed back to the fixation cross when the stimulus ceased. The shapes were counterbalanced across participants. Each inspiratory load was followed by an unloaded period of between 30-60 seconds that was indicated by a third visual cue. The use of relatively long breathlessness stimuli was chosen to maximize the emotional responses associated with anticipation of breathlessness. Participants rated their respiratory intensity and unpleasantness after each stimulus. Visual stimuli were generated and presented in white on a black background using the Cogent toolbox (www.vislab.ucl.ac.uk/Cogent/) for MatLab (MathWorks Inc., Natick, MA, USA).

In an externally cued delay conditioning paradigm (**Supplementary Fig. 2**), participants learned associations between three visual cues (conditioned stimuli, (CS), either a white square, star or triangle shape presented on a black background) and resistive inspiratory loading that was intermittently applied to induce three different respiratory sensations (unconditioned stimuli, (US) - either breathlessness (strong inspiratory load, approximately - 12cmH_2_O), a mild inspiratory load (approximately -3cmH_2_O) or no inspiratory load). The pairing between the visual cue (CS) and respiratory load (US) was maintained constant for each participant during all 3 experimental sessions but was counterbalanced between participants. Four repeats of each of the mild and strong load, and eight repeats of the unloaded condition were performed. Immediately after each inspiratory load, participants rated the intensity and unpleasantness of their breathing on a horizontal visual analogue scale (VAS) with the anchors ‘no breathlessness’ on the left and ‘severe breathlessness’ on the right for intensity ratings, and ‘not unpleasant’ on the right and ‘extremely unpleasant’ on the left for unpleasantness ratings. There was no correlation between the pressure generated and the subjective ratings reported. The inspiratory loads did not extend to intolerable loads. Bond-Lader mood values of tension-relaxation, sedation-alertness, and discontentment-contentment (21) were obtained immediately following the breathlessness protocol using visual analogue scales (VAS) displayed on the screen and a button box. At the end of the experiment, participants were debriefed using the multidimensional dyspnea profile (MDP) questionnaire (22).

Conscious association between CS and US was confirmed in writing and 4 participants who did not form such associations were excluded from the study. Following the breathlessness protocol, an anaesthetist performed a medical assessment; this included a 20-minute test infusion of remifentanil to ensure tolerance and safety. Participants were allowed at least 5 minutes to get accustomed to the breathing system before the experiment commenced. Throughout the experiment, the partial pressure of expired carbon dioxide (P_ET_CO_2_) was maintained constant (isocapnia). This was achieved by initially increasing P_ET_CO_2_ by +0.3kPa from baseline by adding CO_2_ to the inspired air, and then manually adjusting inspired CO_2_ as necessary (23, 24). P_ET_CO_2_ was maintained constant for each participant, and we did not attempt to maintain equivalent P_ET_CO_2_ between subjects. The partial pressure of expired oxygen (P_ET_O_2_) was kept constant at 20kPa in a similar manner.

**Supplementary Figure 3.**
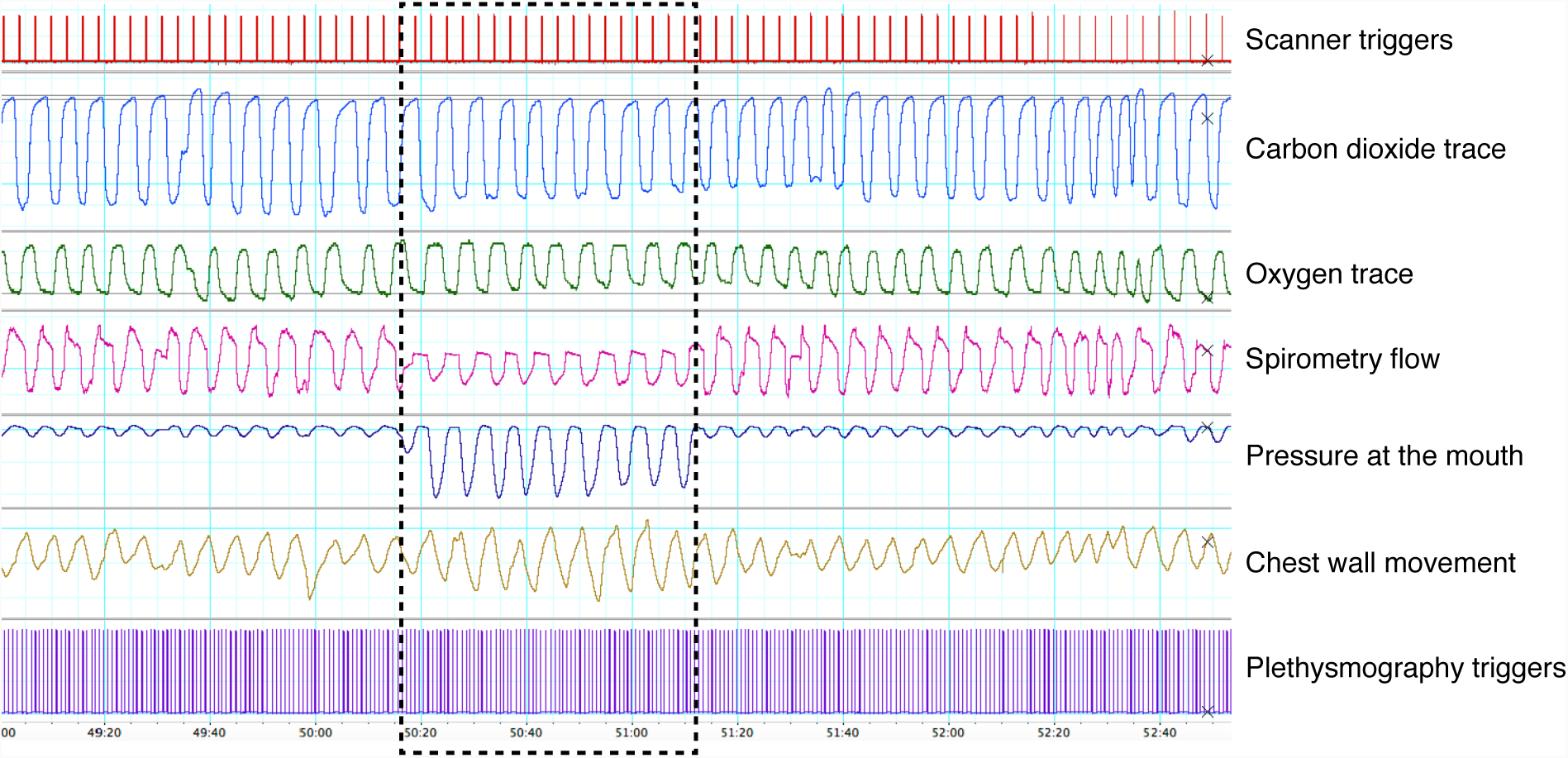
Example trace of the physiological recordings collected throughout the experiment with healthy volunteers. Black dashed box represents an example application of an inspiratory loading stimulus, designed to induce the perception of experimental breathlessness. Scanner triggers: Output from the MR scanner to indicate the beginning of each brain volume measurement. Carbon dioxide trace: Recorded at the mouth, and using small manual additions of 5% Carbon dioxide (in air) mixture this was maintained at +0.3 kPa above baseline. Oxygen trace: Recorded at the mouth, and using small manual additions of a 15% Oxygen (balance nitrogen) mixture this was maintained at approximately 20 kPa. Both end-tidal carbon dioxide and oxygen were manually adjusted on a breath-by-breath basis to minimize fluctuations in these values. Spirometry flow: Ventilatory flow as measured by a spirometer, demonstrating a reduction in flow during the stimulus. Pressure at the mouth: Pressure trace measured at the mouth, demonstrating a marked increase in inspiratory pressure during the application of the stimulus. Chest wall movement: Measured via a respiratory bellows belt around the chest, demonstrating a small increase in tidal volume during the stimulus. Plethysmography triggers: Cardiac cycle peaks measured via pulse oximetry, demonstrating no observable difference in heart rate during the application of the stimulus.

### Physiological recordings

Arterial oxygen saturation and heart rate were monitored continuously, and non-invasive blood pressure was recorded between each scan (In Vivo Research, Orlando, FL, USA). P_ET_CO_2_ and P_ET_O_2_ were determined using rapidly responding gas analysers (ADinstruments ML206) and continuously displayed and recorded with a data acquisition device (PowerLab 8, ADInstruments Ltd, Oxford, United Kingdom) connected to a laptop computer using dedicated software (Chart 5, ADInstruments Ltd, Oxford, United Kingdom)). Inspiratory gas flow was measured with a pneumotachograph (ADinstruments Spirometer FE141) and a standard flow head. Mouth pressure was measured using a Validyne pressure transducer (Validyne Engineering, 8626 Wilbur Ave Northridge, CA 91324). An example trace of physiological recordings collected throughout an experiment is presented in **Supplementary Fig. 3**.

### FMRI sessions

Two FMRI sessions were performed, one on each of the two days following the initial conditioning session, and consisted of a remifentanil and saline placebo session (counterbalanced). During acquisition of the FMRI scans, participants underwent the breathlessness protocol as described above (identical to the initial session). Additional structural scans, field maps and measures of cerebral blood flow were obtained (described in detail below).

### Drug infusion

An intravenous infusion of either remifentanil or saline placebo was delivered in a counterbalanced, randomised and double-blind fashion. The remifentanil dose was modelled on a concentration of 0.7 ng/ml in the brain, achieved using a target-controlled infusion (TCI) pump (Graseby 3500 TCI incorporating Diprisor, SIMS Graseby Ltd, Watford, UK). The TCI pump was pre-programmed with the three-compartment pharmacokinetic model of remifentanil (25, 26). This model controls the infusion rate of remifentanil to achieve and maintain the desired effect site concentration based on the participant’s gender, age, weight and height. The total duration of the infusion was 45 minutes, which allowed for a 10-minute ramp-up period to reach the desired effect site concentration. All participants fasted for 6 hours before each visit and were monitored for an hour after termination of the infusion. Both study participants and study experimenters were blinded to the order of drug administration, with only the administering anaesthetist and the MRI scanner operator being aware of drug condition.

We used remifentanil as a model opioid because its pharmacokinetics and pharmacodynamics are ideal for a mechanistic volunteer study such as this. It is a synthetic μ-opioid agonist with a rapid onset of action, a context sensitive half-life of 3-4 min, and an elimination half-life of approximately 8-10 minutes. This means that the drug has extremely rapid onset and offset, and when combined with a target-controlled infusion, drug levels can be easily manipulated in a short time frame. Due to its rapid onset and offset, remifentanil needs to be administered intravenously, and when this infusion is combined with a pharmacokinetic model of its action, it is possible to manipulate plasma and effect site (brain) concentrations in a predictable and consistent manner. Using a target-controlled infusion allows a constant drug effect to be maintained throughout the experiment.

In terms of volunteer safety, if adverse effects develop, the infusion can be terminated with the knowledge that the drug will wear off within minutes. We chose to deliver an effect site concentration of 0.7ng/ml based upon our extensive clinical and experimental experience with this drug (27-34). Although direct comparison of equivalence with other opioids (e.g. oral morphine) is difficult, we chose a dose that is at the lower end of efficacy for the treatment of acute pain, but which has previously been shown to have effects on respiration (29) and pain suppression (27). We would estimate that the effect site concentration of 0.7ng/ml would represent the analgesia offered by 4-7 mg oral (or 2.0-3.5 mg intravenous) morphine used to treat low-moderate pain. Remifentanil is ultra-short acting, and this means that when given as a bolus the effects would vary within the scanning session, making it difficult to dissociate primary drug effect from secondary effects such as raised P_ET_CO_2_. For this reason, comparison with studies employing bolus doses of remifentanil (35) is difficult.

### MRI data acquisition

MRI data were acquired on a 3 Tesla Siemens Trio scanner using a 32-channel head coil. A whole-brain gradient echo, echo-planar-imaging (EPI) sequence (TR = 3000 ms, TE = 30 ms, field of view: 192×192 mm, voxel size 3×3×3 mm, 45 slices, 380 volumes) was used for functional scans. Fieldmaps were obtained using a symmetric-asymmetric spin-echo sequence before the baseline functional scans (30 ms echo time, 0.5494 ms dwell time, field of view and matrix identical to EPI) and were used to correct for magnetic field inhomogeneities. A T1-weighted structural (MPRAGE sequence, repetition time (TR) = 1720 ms, echo time (TE) = 4.68 ms, flip angle 8°, voxel size 1×1×1 mm) image was acquired for functional image registration.

### FMRI image pre-processing

FMRI data processing was carried out using FEAT (FMRI Expert Analysis Tool) Version 6.00, part of FSL (FMRIB’s Software Library, www.fmrib.ox.ac.uk/fsl) using a whole-brain approach. Pre-processing of the data was performed with MCFLIRT motion correction (36), non-brain removal using BET (37), spatial smoothing using a full-width-half-maximum Gaussian kernel of 5 mm, high pass temporal filtering (Gaussian-weighted least-squares straight line fitting, with sigma = 75.0 s) with 150 seconds cut-off. Registration to high resolution structural was carried out using boundary-based-registration (BBR; FSL software library), registration from high resolution structural to standard space was then further refined using nonlinear registration (FNIRT; FSL software library).

The FMRI scans were corrected for motion, scanner and cerebrospinal fluid artefacts using independent component analysis (ICA) denoising (38). BOLD images were corrected for physiological noise with physiological noise modelling integrated in FEAT (RETROICOR) (39, 40). To ensure noise was not reintroduced through the combination of ICA denoising and RETROICOR, the following stepwise process was followed:

1. ICA was used to decompose FMRI data into different spatial and temporal components using FSL’s MELODIC (Multivariate Exploratory Linear Optimised Decomposition into Independent Components) using automatic dimensionality estimation. This is a data-driven approach to ICA dimensionality estimation, which allows MELODIC to choose the number of components into which the FMRI data is decomposed. This means that different FMRI scans may be decomposed into a different number of components. The noise components (classified as movement, cerebral spinal fluid (CSF) and scanner artefact) were then identified based on spatial location of signal and signal frequency (41, 42), and manually removed from the 4D FMRI data, this is referred to as the “ICA denoised data”.
2. The pulse oximetry and respiratory flow measurements allowed determination of cardiac and respiratory phase with each acquired slice of each FMRI image volume. This assigned phase is then entered into a low-order Fourier expansion (40, 43) to derive time course regressors (2 cardiac, 2 respiratory and 1 interaction regressor). These regressors explain potential signal changes associated with cardiac and respiratory (or interacting) processes. These regressors were regressed against the ICA denoised data using FEAT’s Physiological Noise Modelling (PNM) tool (40, 43). The residuals from this regression were subtracted from the ICA denoised data. This is referred to as the “PNM signal”.
3. To combine the ICA denoising with PNM it is necessary to remove any overlap between the two. Therefore, the PNM signal was also run through ICA denoising (41, 42) and the resulting denoised PNM signal was removed from the ICA denoised data to give the final denoised data used in the analysis.

### First level FMRI analysis

Linear models were used to describe the data. First level analysis used a general linear modelling (GLM) approach with multiple explanatory variables (EVs). Individual subject contrasts were generated for mild and strong anticipation periods from the beginning of the symbol presentation until the onset of the inspiratory resistive loading. ‘On’ loading periods for each of the mild and strong stimuli were then constructed from the onset of the resistance stimulus until the end of the loading period. Unloaded periods were modelled for the duration of the unloaded stimulus. Periods when participants rated their preceding sensations and the 5-second periods after each respiratory manipulation (fixation cross; presented to give participants a chance to recover before giving their subjective ratings) were modelled as regressors of no interest. P_ET_CO_2_ was entered as a separate EV in order to account for residual fluctuations in CO_2_ that could affect BOLD signal (24).

To account for possible changes in the haemodynamic response function (HRF), including slice-timing delays, external timing files were convolved with an optimal basis set of three waveforms (FLOBS: FMRIB’s Linear Optimal Basis Sets, default FLOBS supplied in FSL (44)), instead of the standard gamma waveform. The second and third FLOBS waveforms, which model the temporal and dispersion derivatives, were orthogonalised to the first waveform, of which the parameter estimate was then passed up to the higher level to be used in group analysis.

### Higher level FMRI analysis

Using this brain imaging data, we aimed to determine how the extent of an individual’s response to remifentanil related to their brain activity at baseline (i.e. during infusion of 0.9% saline, the placebo condition). This analysis was designed to explore where the brain responses to breathlessness and anticipation during the saline condition were indicative of the subsequent magnitude of remifentanil-induced relief of breathlessness. A higher-level mixed effects analysis was conducted for the saline condition only. A principal component analysis (PCA; MATLAB 2013a) was performed on the opioid ‘response’ cluster from the hierarchical cluster analysis. The resulting individual scores were included within a group fMRI analysis in the saline condition only, using a general linear model (Z > 2.3, whole brain corrected p < 0.05). FMRIB’s Local Analysis of Mixed Effects (FLAME 1+2 (44)) was used with automatic outlier de-weighting. Analysis was corrected for multiple comparisons across the whole brain, a cluster threshold of >2.3 and a corrected cluster significance threshold of p = 0.05 were used.

This analysis differs from that presented in Hayen et al. (2), which examined group mean differences rather than exploring potential mechanisms underlying inter-individual variability in response. The brain imaging data collected during remifentanil administration was not analysed and is not presented in this manuscript.

## Hierarchical cluster threshold techniques

For both hierarchical cluster analyses performed in this manuscript, the elbow method and visual inspection of dendrogram structure were utilized for cluster thresholding and separation. The elbow method is a validated clustering technique in which the percentage of explained variance is described as a function of the number of clusters. Considering the variable set as initially one large cluster, the algorithm then divides the variables into increasing numbers of clusters. With each additional cluster, the percentage of explained variance is expected to increase. While initially this increase is sharp, after a certain number of clusters the gain will become marginal. When this relationship is plotted, as the sum of intra-cluster distance against cluster number, the point at which additional clusters add only marginally to the explained variance can be seen as a sharp bend or elbow in the graph. The number of clusters corresponding to this elbow point is thus the number of most statistically distinct clusters in the dendrogram.

The elbow method is, however, fundamentally limited by the stability of the variables within a cluster. When situations arise where one or more variable sits at the border of two clusters the method can give unreliable results. In the current investigation, the elbow method was trialed and assessed for test-retest reliability (a marker of cluster stability), across 10 trials. Although there was clear variance across the trials in both the healthy and COPD datasets, the overall trend indicated 2 or 3 major clusters within the healthy population, and 3 major clusters within the COPD population (**Supplementary Fig. 4**). Whilst additional modeling techniques such as confirmatory factor analysis (CFA) could then be employed to validate clusters and sub-clusters, there are insufficient sample sizes in these datasets for a valid CFA to be conducted. Therefore, in this work, the elbow method was applied and followed by visual inspection of the dendrogram structure.

**Supplementary Figure 4.**
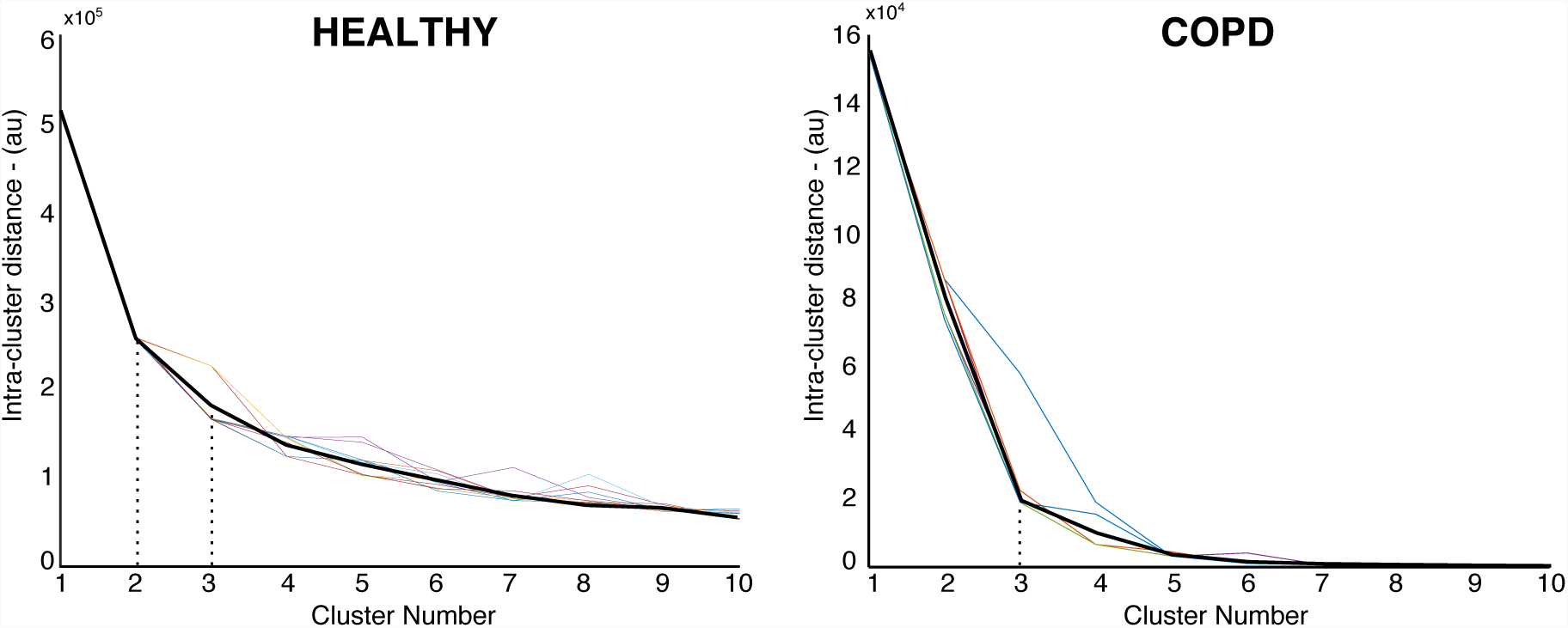
Elbow plots - where variability within a cluster (y-axis, au - arbitrary units) is plotted as a function of the number of clusters (x-axis) for both the healthy (left) and COPD (right) datasets. The point at which intra-cluster distance is no longer sharply decreased by adding further clusters can be visualised as an “elbow” in the plot. Each coloured line represents one trial (10 trials total), while the thicker black line represents the average of all trials. Dotted lines highlight the “elbow joint” of the graph. The elbow plot for the healthy population contains two possible joint points, one at 2 clusters and one at 3 clusters, while the average elbow line clearly bends at 3 clusters within the COPD population dataset.

## RESULTS

The inspiratory resistive loads used in the healthy control study induced a sensation of breathing intensity and unpleasantness. Compared to unloaded breathing, the mild and strong inspiratory loads significantly increased breathlessness intensity (mean VAS%; unloaded vs. mild loading: 12 vs. 32%; unloaded vs. strong loading: 12 vs. 71%) and breathlessness unpleasantness (mean VAS%; unloaded vs. mild loading: 10 vs. 25%; unloaded vs. strong loading: 10 vs. 61%)). Furthermore, the inspiratory resistive load increased the work-effort and air hunger sensations (**Supplementary Fig. 5**).

**Supplementary Figure 5.**
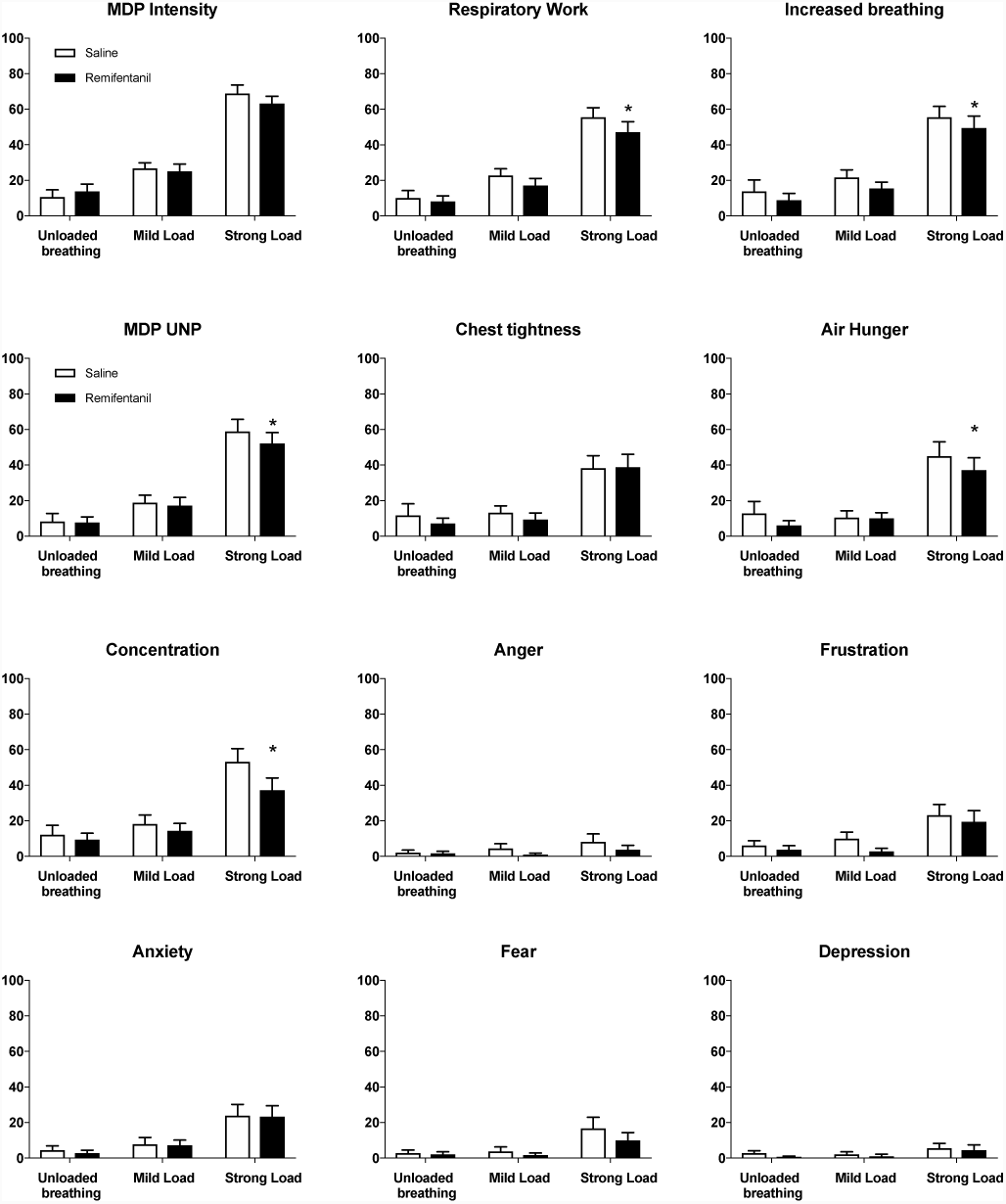
Sensory and affective dimensions of dyspnea, as measured with the multidimensional dyspnea profile (MDP), during saline and remifentanil. Mean±SEM. *significant difference between saline and remifentanil condition at p<0.05.

